# PhyloAln: a convenient reference-based tool to align sequences and high-throughput reads for phylogeny and evolution in the omic era

**DOI:** 10.1101/2024.02.08.579425

**Authors:** Yu-Hao Huang, Yi-Fei Sun, Hao Li, Hao-Sen Li, Hong Pang

## Abstract

The current trend in phylogenetic and evolutionary analyses predominantly relies on omic data. However, traditional methods typically involve intricate and time-consuming procedures prior to core analyses. These procedures encompass assembly from high-throughput reads, decontamination, gene prediction, homology search, orthology assignment, multiple alignment, and matrix trimming. Such processes significantly impede the efficiency of research when dealing with extensive datasets. In this study, we present PhyloAln, a convenient reference-based tool capable of directly aligning high-throughput reads or complete sequences with existing alignments as reference for phylogenetic and evolutionary analyses. Through testing with both simulated and authentic datasets, PhyloAln demonstrates consistently robust performance in terms of alignment completeness and identity when compared to other reference-based tools. Additionally, we validate the tool’s adeptness in managing foreign and cross-contamination issues prevalent in sequencing data, which are often overlooked by other tools. Moreover, we showcase the broad applicability of PhyloAln by generating alignments and reconstructing phylogenies from transcriptomes of ladybird beetles, plastid genes of peppers, and ultraconserved elements of turtles. These results underscore the versatility of our tool. Leveraging these advantages, PhyloAln stands poised to expedite phylogenetic and evolutionary analyses in the omic era. The tool is accessible at https://github.com/huangyh45/PhyloAln.

## Introduction

Phylogenetic analyses elucidate relationships and evolutionary histories across various species (Misof, et al. 2014; Zeng, et al. 2014; Prum, et al. 2015), strains (Lemieux, et al. 2021), individuals (Prüfer, et al. 2014; Sun, et al. 2023), cells (Coorens, et al. 2021) and genes (Li, et al. 2022). Furthermore, phylogenetic trees play a pivotal role in downstream applications, including ancestral state reconstruction, testing evolutionary hypotheses, conducting phylogenetic comparative and diversity analyses, and consequently, they are widely used in research of evolutionary biology, conservation biology, earth history, and ecology (Mitchell, et al. 2015; Lu, et al. 2018; Liang, et al. 2023; Siqueira, et al. 2023). The advent of high-throughput sequencing technology has facilitated the generation of extensive omic datasets, encompassing genomes, transcriptomes, amplicons, and plastomes. These datasets contribute significantly to the establishment of robust phylogenies across the tree of life. Examples include the elucidation of phylogenetic relationships among major lineages in insects (Misof, et al. 2014), the evolutionary history of the living birds (Prum, et al. 2015), highly supported genomic phylogeny of living primates (Shao, et al. 2023), the position of the eukaryotes within Asgard archaea (Eme, et al. 2023), and the deep phylogeny of angiosperms (Zeng, et al. 2014). In addition, alignments conforming to standards for phylogenetic analyses find utility in various evolutionary studies beyond tree reconstruction. These alignments can be employed in analyses such as the detection of selection pressure and site substitutions (Wang, et al. 2020; Peng, et al. 2023; Shao, et al. 2023).

However, phylogenetic analyses relying on omic data, such as phylogenomic and phylotranscriptomic analyses, consistently involve intricate and time-consuming preparatory steps to generate sequence alignment matrices. Traditional preparation encompasses multiple procedures, including data assembly from raw reads, contaminant identification, gene prediction, orthology assignment, sequence alignment, and site trimming (Kapli, et al. 2020). Protein datasets derived from assembled omic data often serve as the basis for phylogenetic inferences. These inferences primarily rely on grouping proteins through *de novo* orthology assignments, exemplified by methods such as OrthoMCL (Li, et al. 2003) and OrthoFinder (Emms and Kelly 2019). Moreover, omic data provide support for phylogenetic analyses employing sequences beyond nuclear protein-coding genes. Examples include the utilization of ultraconserved elements (UCEs) (Faircloth, et al. 2012), ribosomal genes, mitochondrial sequences and plastid sequences (Patwardhan, et al. 2014).

To better and more efficiently harness omic data, strategies have been developed for generating target sequences or alignments for phylogeny based on reference alignments. Well-established ortholog databases, such as OrthoDB (Kuznetsov, et al. 2023) and OMA (Altenhoff, et al. 2023), offer extensive references for these reference-based methods, exemplified by tools like Orthograph (Petersen, et al. 2017) and Read2Tree (Dylus, et al. 2024). Orthograph, primarily designed for phylotranscriptomics, employs a best reciprocal hit strategy to search transcriptomic sequences based on the globally best-matching cluster of orthologous genes (Petersen, et al. 2017). However, Orthograph is limited to assembled sequences and its output includes only target sequences without alignments. In contrast, Read2Tree is a novel tool utilizing an assembly-free strategy that directly maps raw sequencing reads into groups of corresponding genes, generating consensus sequences, alignments, and phylogenetic trees (Dylus, et al. 2024). Nevertheless, Read2Tree is tailored for raw reads and features relatively inflexible processes, allowing only unaligned reference sequences and FASTQ read files as input. Additionally, these tools do not account for common foreign and cross-contamination, such as those from parasites, endosymbionts, or other species in the same batch, which can potentially impact phylogenetic analyses (Lusk 2014; Borner and Burmester 2017; Simion, et al. 2018). Moreover, these tools may not adequately handle sequences distinct from nuclear protein-coding genes, including protein-coding sequences with nonstandard genetic codes (e.g., mitochondrial and plastid genes), non-protein-coding sequences (e.g., UCEs and ribosomal genes), and translated amino acid sequences.

In an effort to better address the requirements of phylogenetic and evolutionary analyses, this article introduces PhyloAln—a reference-based alignment tool designed for phylogeny and evolution. PhyloAln has the capability to directly map not only raw reads but also assembled or translated sequences onto reference alignments. This feature makes it applicable to various omic data types, eliminating the need for the complex preparation involved in traditional methods, such as data assembly, gene prediction, orthology assignment, and sequence alignment. Furthermore, PhyloAln is equipped to identify and eliminate foreign and cross contamination within the generated alignments—an aspect often overlooked in other reference-based methods. This capability enhances the quality of alignments for subsequent analyses. To evaluate its performance, we subjected PhyloAln to testing with simulated datasets, including alignments of single-copy orthologous genes spanning the tree of life and contaminated transcriptomes of fruit flies. Real datasets, consisting of alignments from transcriptomes of ladybird beetles, plastid genomes of peppers, and ultraconserved elements of turtles, were also used for assessment. Comparative analyses with results from Read2Tree and Orthograph were conducted to gauge the effectiveness of PhyloAln. The assessments show that, PhyloAln exhibits a relatively high accuracy in aligned sites and downstream phylogenetic analyses and broad availability in different data.

## Results

To streamline the intricate and time-consuming process of preparing multiple sequence alignments (MSAs) using omic data for phylogenetic and other evolutionary analyses, we introduce PhyloAln. It is a command line tool based on Python framework and designed to map not only assembled sequences but also raw reads onto reference alignments, accommodating various data types (Figure 1). The workflow begins with the preprocessing of reference alignments and target sequences or reads, incorporating optional reverse complement and translation based on user-defined modes. Subsequently, the processed sequences and reads undergo a search against the reference alignments using HMMER3 software (Mistry, et al. 2013), and the mapped sequences are aligned to the reference coordinates derived from HMMER3 search results. These mapped sequences undergo optional assembly, foreign and cross-contamination removal, and consensus steps. Finally, the resulting mapped sequences are included in the output DNA and protein alignments, providing new alignment files suitable for downstream phylogenetic analyses. Further details for each step are outlined in the Methods section.

**Figure 1.**
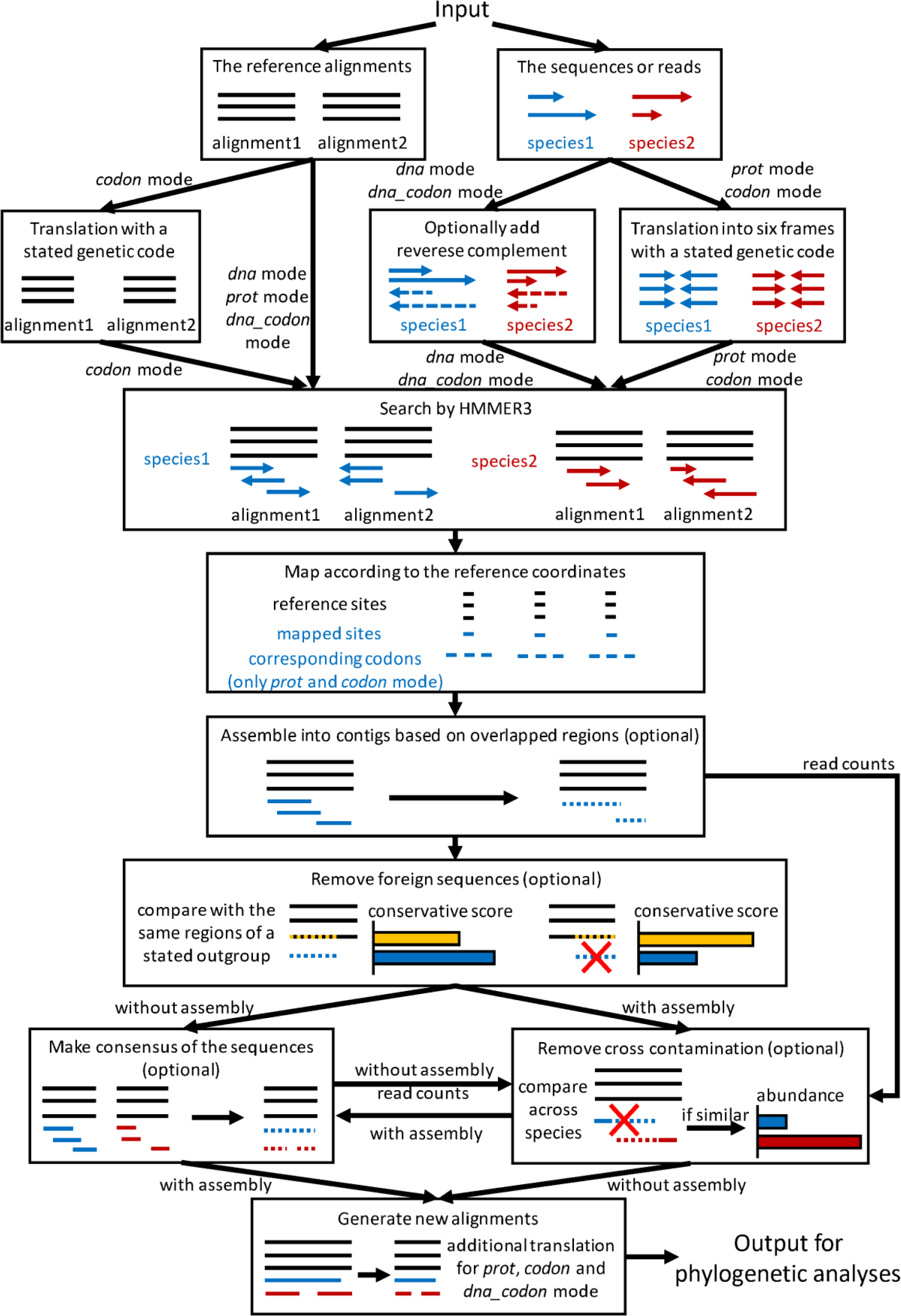
Overview of the strategy and technological processes employed by PhyloAln.

### Performance on simulated data across life of tree with different coverages

We conducted a comparative analysis of the alignment performance between PhyloAln and another reference-based tool, Read2Tree (Dylus, et al. 2024), utilizing simulated reads encompassing different omic data (genomes and transcriptomes), sequencing technologies (Illumina, PacBio, and Nanopore), coverages (2X, 5X, 10X, 20X, 30X, and complete assembly sequences), and species across the life of the tree (*A. thaliana*, *H. sapiens* and *D. melanogaster*; Figure 2A). In the case of short Illumina reads, PhyloAln consistently outperformed Read2Tree, yielding more complete and identical alignments of single-copy orthologous genes (average completeness: 51.61-90.44%, average identity: 83.19-97.82%, of all genes) in the three target species, with the alignments based on *de novo* orthology assignment using gene predictions from the assembly sequences as reference (Figure 2B). Read2Tree exhibited lower performance (average completeness: 0-19.55%, average identity: 0-35.25%). PhyloAln demonstrated comparable or even shorter runtime across most scenarios (1.13-175.25 min, except for 1717.44 and 2728.22 min for the extremely large genomic data of *H. sapiens* with the coverage of 20X and 30X, respectively, using the slower low-memory strategy) in comparison to Read2Tree (1.70-628.94 min) for aligning Illumina short reads (Figure 2C, Table S2). Moreover, in contrast to Read2Tree, which works only with reads, PhyloAln efficiently mapped assembly sequences directly into alignments with high completeness (71.12-97.99%) and identity (74.99-94.88%), requiring minimal time (0.44-5.34 min, except 158.52 for *H. sapiens* genome using the low-memory strategy). For long PacBio and Nanopore reads, PhyloAln demonstrated the capability to generate alignments with relatively high completeness (75.95-96.37%) and moderate identity (45.49-84.52%) (Figures S1 and S2).

**Figure 2.**
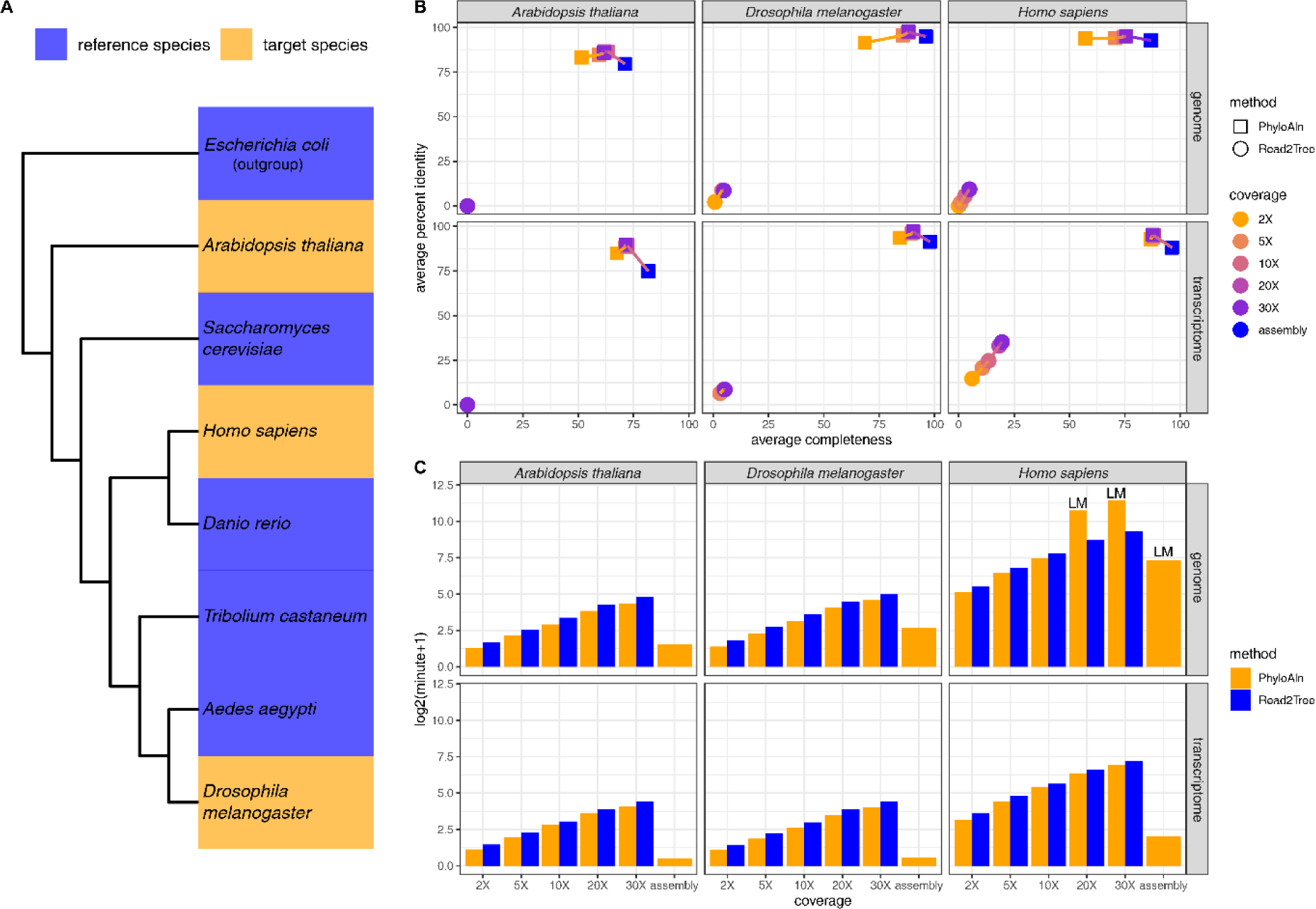
Performance test of PhyloAln and Read2Tree on the simulated dataset across the tree of life. A) Design for the dataset. B) Average completeness and percent identity of the alignments generated by PhyloAln and Read2Tree using simulated Illumina reads or original assemblies of genomes and transcriptomes from three target species with varying coverages, with the MAFFT alignments based on *de novo* gene orthology assignment by OrthoFinder as reference. C) Running time of PhyloAln and Read2Tree using simulated Illumina reads or original assemblies of genomes and transcriptomes from three target species with different coverages. LM: using a slower low-memory strategy to prepare the sequences and reads because of the large size of the data.

### Ability to remove foreign and cross contamination

We evaluated PhyloAln’s capability to eliminate foreign and cross contamination, common issues in sequencing data (Lusk 2014; Borner and Burmester 2017; Simion, et al. 2018) often overlooked by other reference-based tools. This assessment was conducted using a simulated dataset of contaminated fruit fly transcriptomes (Figure 3A). Additionally, we conducted a comparative analysis of the overall performance on the alignments generated by PhyloAln and Read2Tree. Regarding the completeness and identity of the alignments, both PhyloAln with and without the assembly step based on the reference coordinates demonstrated strong performance. The assembly step exhibited slightly superior alignments (average completeness: 96.70-98.42%, average identity: 98.70-99.43% with the assembly step versus average completeness: 95.64-98.17%, average identity: 98.81-99.26% without the assembly step; Figure 3B). Read2Tree also exhibited good performance on the aligned sequences of *D. melanogaster* and *D. simulans* (average completeness: 99.34-99.65%, average identity: 99.79-99.80%). However, its performance was relatively weaker on the sequences of *D. willistoni* (average completeness: 78.47%, average identity: 80.92%).

**Figure 3.**
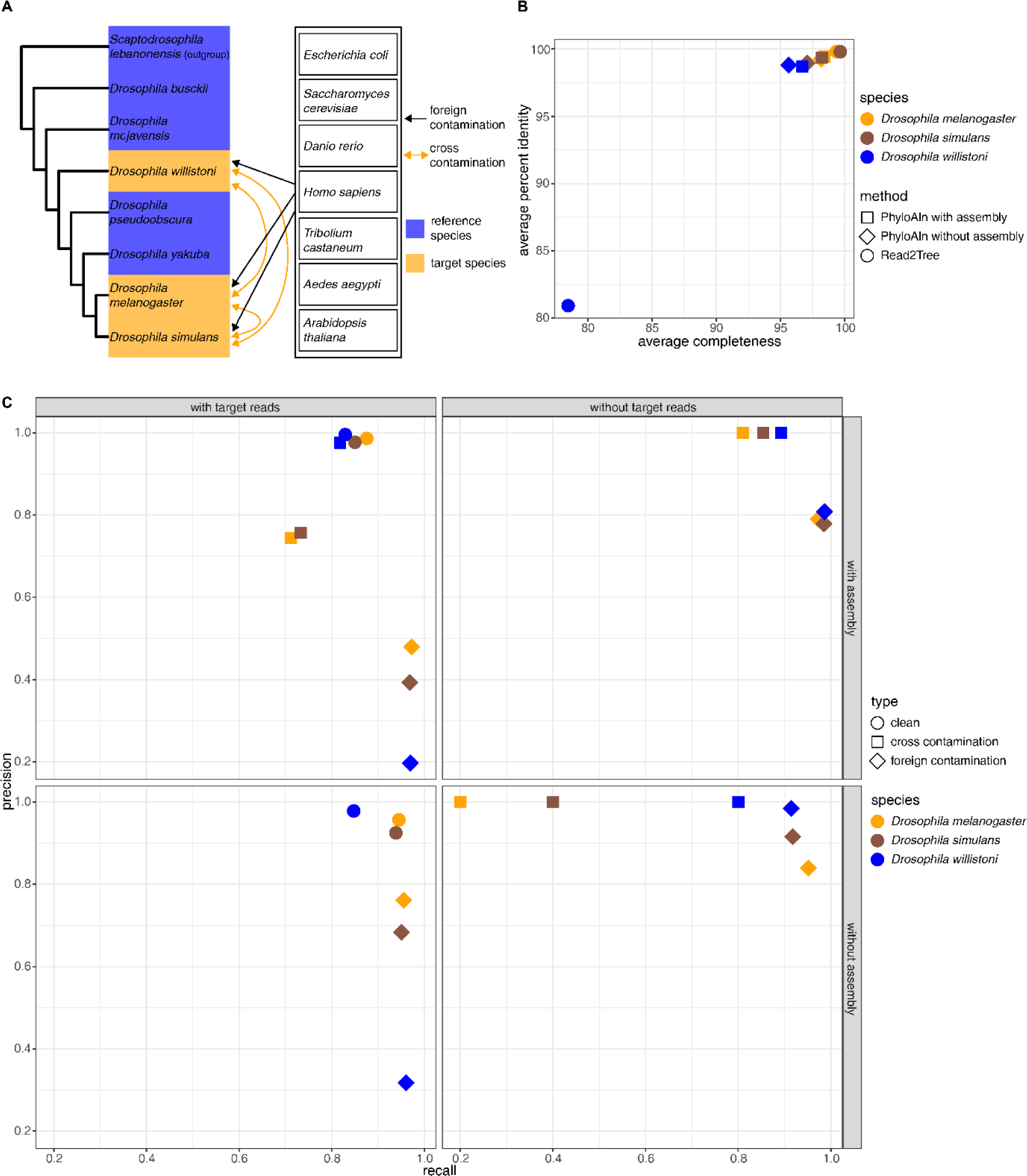
Performance test of PhyloAln and Read2Tree on the simulated dataset of contaminated fruit fly transcriptomes. A) Design for the dataset. B) Average completeness and percent identity of the alignments generated by PhyloAln and Read2Tree using simulated reads of contaminated transcriptomes from three target fruit fly species, with the MAFFT alignments based on de novo gene orthology assignment by OrthoFinder as reference. C) Precision and recall of PhyloAln to align the reads from the corresponding target species (clean reads) and remove the reads of foreign contamination and cross-contamination with or without the assembly step and the target reads.

To assess PhyloAln’s decontamination efficacy, we quantified the precision and recall of aligned reads (clean), as well as reads removed during the foreign and cross contamination removal steps, categorized by their species sources. PhyloAln consistently demonstrated high precision (0.925-0.996) and recall (0.829-0.945) in aligning clean reads from the target species, both with and without the assembly step (Figure 3C). In orthologous genes with target reads in the data, PhyloAln successfully removed 95.07-97.27% of foreign contamination with a precision of 0.197-0.761, irrespective of the assembly step. It also eliminated 71.21-81.85% of cross contamination with a precision of 0.744-0.976 when the assembly step was applied. Notably, PhyloAln without the assembly step did not detect any cross contamination reads, likely due to their coverage by the consensus with a substantial number of clean reads, resulting in minimal impact on the output alignments (Figure 3B). In the five orthologous genes where we retained only contamination reads by removing target reads, PhyloAln, with or without the assembly step, successfully removed 91.44-98.63% of foreign contamination reads with a precision of 0.779-0.984. However, the assembly step detected 80.97-89.27% of cross contamination, compared to 20-80% detected without this step.

Upon observing relatively low precision in foreign contamination detection using PhyloAln, with or without the assembly step, in orthologous genes with target reads, we examined the alignments generated by the consensus specifically for reads detected as foreign contamination but originating from the target species. We observed that these mapped reads in the alignments exhibited poor alignment quality, mostly with lower percent identity compared to the output alignments of PhyloAln (32-35 out of 41 alignments with retained target reads; Figures S3 and S4). This suggests that these poorly aligned reads are primarily attributed to paralogous genes, randomly similar sequences, or hypervariable regions, making limited contributions to a robust phylogeny. The phylogenetic trees reconstructed from these poor alignments, which exhibited incorrect positions for the three target species, further support our inference (Figure S5).

### Performance on the real dataset of the ladybird beetle transcriptomes

Furthermore, we evaluated the performance of PhyloAln using both reads and assemblies, Read2Tree employing reads, and Orthograph (Petersen, et al. 2017) utilizing assemblies on a real dataset of ladybird beetle transcriptomes (Li, Tang, et al. 2021). PhyloAln effectively mapped reads and assemblies to the reference alignments, consistently achieving high completeness (94.63-99.58% with reads and 92.61-99.65% with assemblies) and identity (96.26-99.13% with reads and 92.63-99.27% with assemblies). In contrast, Read2Tree results exhibited completeness ranging from 66.29% to 99.96% and identity ranging from 82.35% to 99.09%, while Orthograph achieved sequence searches with completeness between 98.95-100% and identity between 70.54-98.85% (Figure 4A). The subsequent phylogenetic tree reconstructed from the alignments revealed that all nodes, in both the PhyloAln-based trees using reads or assemblies, were identical and highly supported, with a UFBoot value of 100 (Figure 4B). The reported monophyletic clades in ladybird beetles, such as Microweiseinae, Coccinellinae, Coccinellini, Epilachnini, ABDHP Clade, CSPS Clade, Scymnini, mycophagous group, and four clades (A-D) in Coccinellini (Che, et al. 2021; Li, Tang, et al. 2021; Nattier, et al. 2021; Tomaszewska, et al. 2021), were all supported in the PhyloAln-based phylogenetic trees (Figure 4C). Although the topologies of the phylogeny reconstructed based on the alignments generated by *de novo* gene predictions and orthology assignments by OrthoFinder (Emms and Kelly 2019), PhyloAln using reads or assemblies, Read2Tree using reads, and Orthograph using assemblies differed at a few nodes (Figures S6-S10), the unique nodes in the PhyloAln-based trees occurred at those with relatively low support UFBoot values of 65-77 in at least one phylogenetic tree using other methods (Figures 4B, S6-S10). Furthermore, the positions of *Megalocaria dilatata* and *Micraspis discolor* in the trees using PhyloAln, conflicting with other methods, are consistent with the reported Coccinellini phylogeny with comprehensive species samples (Nattier, et al. 2021; Tomaszewska, et al. 2021). All these findings suggest potentially more robust trees using PhyloAln.

**Figure 4.**
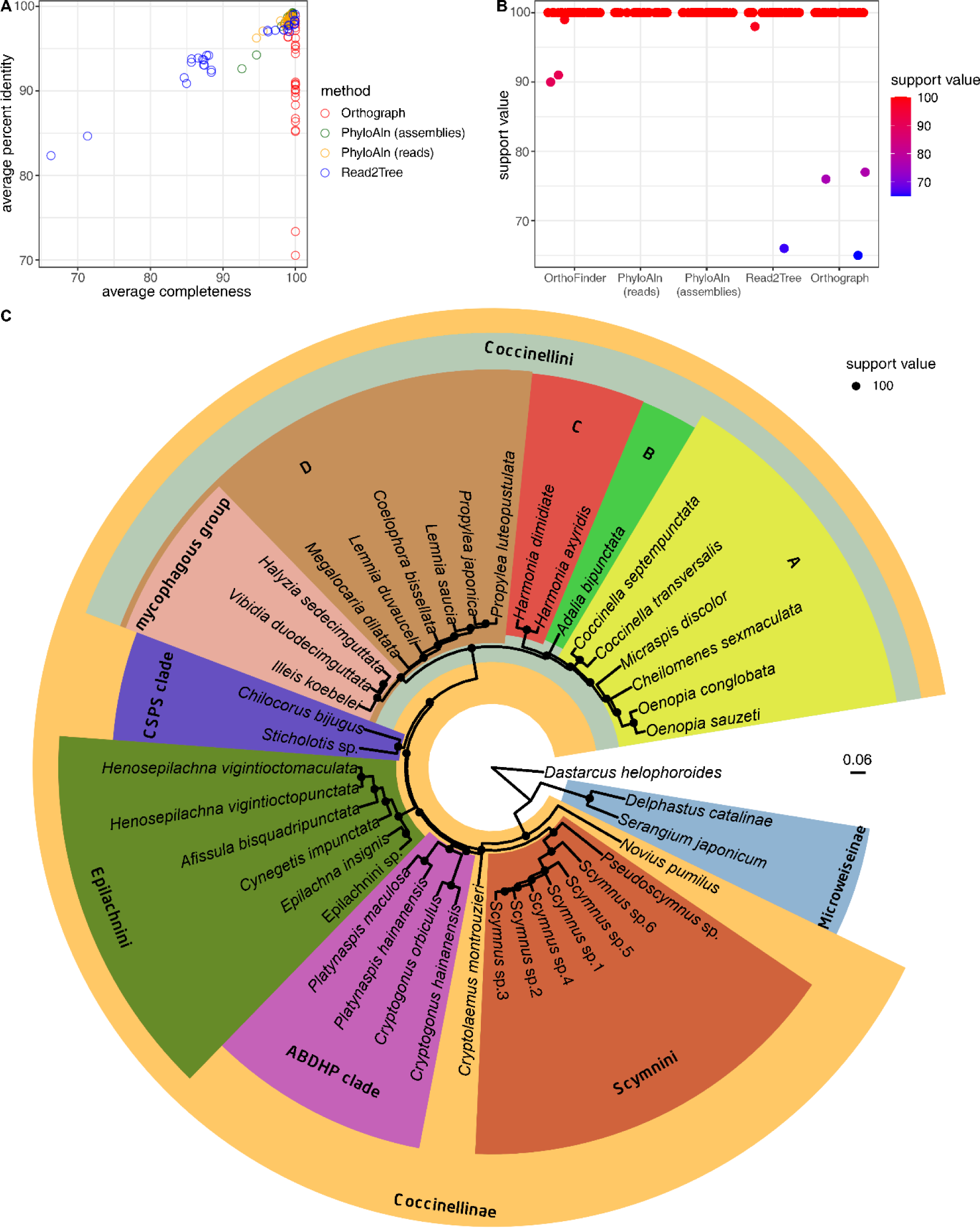
Performance test of PhyloAln, Read2Tree, and Orthograph on the real dataset of ladybird beetle transcriptomes. A) Average completeness and percent identity of the alignments generated by PhyloAln using the reads or assemblies, Read2Tree using the reads, and Orthograph using the assemblies from the 30 target ladybird transcriptomes, with the MAFFT alignments based on *de novo* gene predictions and orthology assignment by OrthoFinder as reference. B) T The ultrafast bootstrap (UFBoot) support values of all nodes in the phylogeny reconstructed by IQ-TREE based on the alignments generated by de novo gene predictions and orthology assignments by OrthoFinder, PhyloAln using the reads or the assemblies, Read2Tree using the reads, and Orthograph using the assemblies. C) The robust phylogenetic tree of the ladybird transcriptomes reconstructed by IQ-TREE based on the alignments generated by PhyloAln using the reads, with the same topology as those produced through PhyloAln using the assemblies. The reported monophyletic clades in the ladybird beetles (Che, et al. 2021; Li, Tang, et al. 2021; Nattier, et al. 2021; Tomaszewska, et al. 2021) include Microweiseinae, Coccinellinae, Coccinellini, Epilachnini, ABDHP Clade, CSPS Clade, Scymnini, mycophagous group and four clades (A-D) in Coccinellini.

### Application on data of pepper plastomes

To showcase the utility of PhyloAln in handling genes with non-standard genetic codes, we employed the tool to map reads from pepper plastomes (Simmonds, et al. 2021) into reference alignments and assessed its performance. The resulting alignments from PhyloAln exhibited remarkably high completeness (99.65-99.98%) and identity (99.57-99.99%) when compared to alignments generated from gene predictions of the plastome assemblies as a reference (Figure 5A). The phylogenetic trees reconstructed from alignments produced by PhyloAln and gene predictions of the assemblies displayed nearly identical topologies, with the exception of the position of *Piper caninum*1 (Figure 5B). In the phylogenetic tree generated using PhyloAln, *Piper caninum*1 emerged as the sister sample to *Piper caninum*2, a relationship that appears more reliable considering the belonging to the same species.

**Figure 5.**
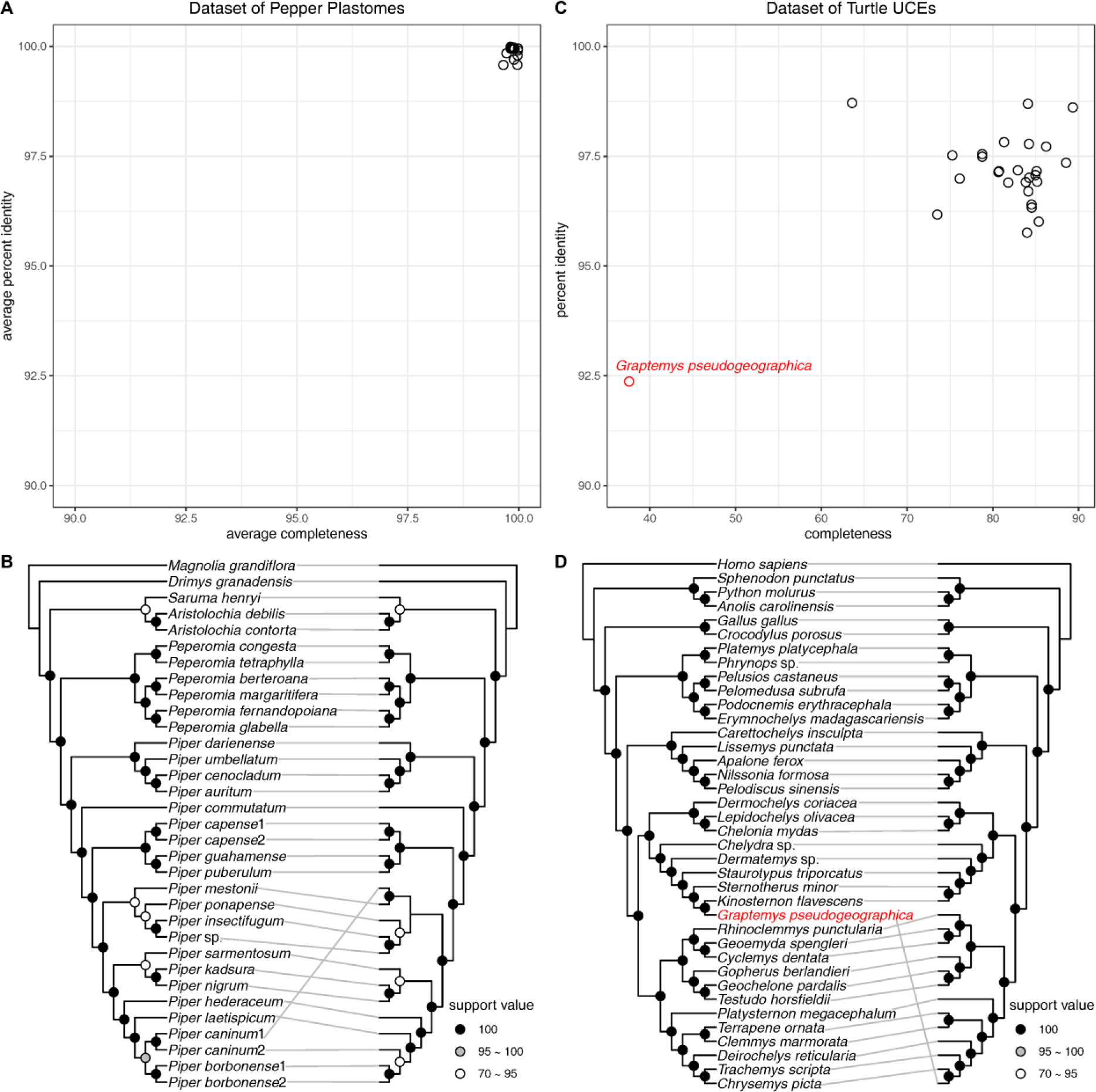
Applications of PhyloAln on the real datasets of pepper plastomes and turtle ultraconserved elements (UCEs). A) Average completeness and percent identity of the alignments generated by PhyloAln using the reads of 17 target pepper plastomes. B) Comparison of the topologies of the phylogenetic trees reconstructed by IQ-TREE based on the alignments generated through PhyloAln using the reads (left) and the gene annotations of the plastome assemblies (right). C) Completeness and percent identity of the UCE result alignments generated by PhyloAln using the reads of UCE sequencing data of 27 target turtles. D) Comparison of the topologies of the phylogenetic trees reconstructed by IQ-TREE based on the alignments generated by PhyloAln using the reads (left) and the downloaded concatenated UCE matrix (right). *Graptemys pseudogeographica*, marked in red, is putatively from different sources of reads and aligned sequences in the downloaded concatenated UCE matrix, thus having different positions in the two phylogenetic trees.

### Application on data of turtle UCEs

To evaluate the applicability of PhyloAln on diverse datasets such as UCEs, a category of DNA genetic markers (Faircloth, et al. 2012), we utilized the tool to map reads to the reference concatenated UCE matrix of turtles (Crawford, et al. 2015). Our tool produced alignments with moderate completeness ranging from 63.58% to 89.32% (except 37.58% for *Graptemys pseudogeographica*) and high identity spanning 95.76-98.71% (except 92.37% for *G. pseudogeographica*; Figure 5C). Phylogenetic trees constructed based on alignments generated by PhyloAln and the published concatenated matrix displayed nearly identical topologies, with the exception of the position of *G. pseudogeographica* (Figure 5D). However, upon scrutinizing the read data of *G. pseudogeographica* downloaded from NCBI, we observed that the assembled sequences from these reads (a total of 3054 sequences with best hits) were primarily closest to *Kinosternon flavescens* (1418 sequences), followed by *Sternotherus minor* (865 sequences) and *Staurotypus triporcatus* (286 sequences), which were the three closest species to *G. pseudogeographica* in our PhyloAln tree (Figures 5D, S11). In contrast, only 12 sequences were closest to the *G. pseudogeographica* sequences in the downloaded supermatrix. Therefore, we concluded that the reads on NCBI and the sequences from the published concatenated matrix originated from different sources, and our PhyloAln tree accurately depicted the correct topology across UCE read sources.

## Discussion

In the current era, an increasing volume of omic data is being sequenced, notably encompassing genomic and transcriptomic datasets rich in gene content, as well as amplicons, mitogenomes, and plastomes, all of which are valuable for phylogenetic analyses. Large-scale initiatives, such as the i5K initiative to sequence 5000 arthropod genomes (Evans, et al. 2013), the Vertebrate Genome Project (Rhie, et al. 2021), and the Darwin Tree of Life Project (The Darwin Tree of Life Project Consortium 2022), contribute to the rapid generation of omic data. With the availability of these extensive omic datasets, there is a growing demand for user-friendly pipelines and tools to efficiently process such data for large-scale phylogenetic and evolutionary analyses (Kapli, et al. 2020). Reference-based alignment has emerged as a novel and viable strategy, leveraging orthology databases (Altenhoff, et al. 2023; Kuznetsov, et al. 2023) and manually *de novo* clustered orthology (Li, et al. 2003; Emms and Kelly 2019) based on part of the high-quality genomes to assign genes from omic data. Here, we introduce PhyloAln, a reference-based alignment tool designed for phylogenetic and evolutionary analyses. PhyloAln offers the capability to map various types of sequences or reads into reference alignments and generate new alignments. This tool provides a more convenient approach for preparing sequence alignments and performing downstream phylogenetic or evolutionary analyses, particularly suited for the developing landscape of high-throughput technologies and the increasing volume of omic data, as compared to traditional methods relying on *de novo* assembly, orthology assignments and sequence alignments (Kapli, et al. 2020).

Based on our tests of completeness and identity, PhyloAln demonstrated effective alignment of genomic and transcriptomic reads and sequences from different sequencing technologies (Illumina, PacBio, and Nanopore), coverages (2X, 5X, 10X, 20X, 30X, and original assembly sequences), and three model organisms across the tree of life, including an insect, a vertebrate, and a plant. Notably, PhyloAln leverages HMMER3 (Mistry, et al. 2013), a profile hidden Markov model-based sequence search tool, ensuring stable performance on conservative sites when mapping target species distantly related to the reference species, spanning families, classes, and even kingdoms. In contrast, Read2Tree, a reference-based tool utilizing expert read aligners, faces challenges in handling target species with divergence larger than hundreds of millions of years from the reference species (Dylus, et al. 2024). Additionally, the performance of *de novo* assembly-based processes is often constrained by low-coverage sequencing data (Liao, et al. 2019; Dylus, et al. 2024). PhyloAln’s reference-based approach allows it to robustly handle reads with low coverage (at least 2X), enabling phylogenetic analyses with low-coverage omic data and, consequently, offering cost-effective and data-abundant sampling capabilities.

In view of the prevalent issues of foreign and cross contamination in omic data (Lusk 2014; Borner and Burmester 2017; Simion, et al. 2018), we have devised methods to address these challenges and incorporated them into PhyloAln. Through tests with simulated fruit fly transcriptomes with contamination, PhyloAln demonstrated the ability to remove a minimum of approximately 90% of foreign contamination, regardless of whether the assembly step was included. It also exhibited the capacity to eliminate at least around 70% of cross contamination with the assembly step, ultimately aligning reads from the target species with high precision ranging from 0.925 to 0.996. Without the assembly step, the ability to detect cross contamination decreased to 20-80%, possibly due to the absence of precise read count information during the assembly step and the potential disturbance of final sequence composition by reads from different contamination sources during the consensus. However, this ability is only essential in unusual conditions where corresponding reads from the target species are absent. In most cases, the substantial number of target reads is likely to offset the impact of relatively few contamination reads in the consensus. Consequently, PhyloAln, with or without the assembly step, can both yield alignments with high completeness and identity. PhyloAln also identifies poorly aligned sequences or reads as foreign contamination and removes them, contributing to a more robust phylogenetic tree compared to other tools.

Unlike other reference-based tools such as Read2Tree (Dylus, et al. 2024) and Orthograph (Petersen, et al. 2017), which either do not consider the contamination issues or necessitate additional decontamination before operation, PhyloAln is specifically designed to handle these common contamination scenarios. This advantage is evident in the stable performance of PhyloAln in alignments compared to Read2Tree, particularly under simulated contamination conditions or real datasets of transcriptomes. Sequences of certain species in alignments generated by Read2Tree exhibited noticeably lower completeness and identity, especially for species less related to the reference species. It is plausible that the read aligner dependency of Read2Tree tends to map reads close to the reference species, resulting in low completeness and attraction to contamination from species closer to the reference sequences, leading to low identity. In contrast, PhyloAln has a distinct advantage in managing contamination scenarios, demonstrating consistently high performance in alignments.

Moreover, the application of PhyloAln to datasets encompassing pepper plastomes and turtle UCEs attests to its broad utility in phylogenetic analyses utilizing diverse omic data from various species groups. Phylogenetic studies commonly leverage not only nuclear protein-coding genes derived from genomes or transcriptomes (Cheon, et al. 2020; Kapli, et al. 2020) but also DNA sequences such as UCEs (Faircloth, et al. 2012), as well as ribosomal genes, mitochondrial sequences and plastid sequences (Patwardhan, et al. 2014). Researchers may encounter a spectrum of data types, including sequenced reads, existing assemblies lacking high-quality gene predictions, predicted gene sequences, and even translated protein sequences. The ability to adeptly process diverse sequence types and accommodate different genetic codes represents an additional advantage of PhyloAln, making it well-suited for a wide array of phylogenetic and evolutionary analyses.

However, it should be noted that PhyloAln is designed as an exclusively reference-based tool, consequently disregarding regions present in the target species but absent or inconsistent in the reference alignments. Therefore, it will consistently miss this information, rendering it unsuitable as a substitute for *de novo* assembly-based methods in cases requiring different standards for phylogenetic analyses. In comparison with the most user-friendly workflows for preparing reference alignments and conducting downstream phylogenetic analyses, we also prioritize the flexibility of use in PhyloAln. This allows the tool to accept previously prepared reference alignments and produce result alignments for user-customized downstream analyses. In addition, we have adopted a tentative approach that does not heavily focus on optimizing the running time and memory usage of PhyloAln. This makes it susceptible to the influence of the number of reference alignments and the sizes of the target sequences/reads. Consequently, this may restrict its ease of use with extremely large or complex datasets in phylogenetic analyses. To address this limitation, PhyloAln could be further developed by optimizing parallel operations and incorporating accessories programmed in faster languages than Python for executing time-consuming steps.

Overall, our reference-based alignment tool, PhyloAln, stands out by offering a stable and direct mapping capability for sequences or reads from diverse omic data to reference alignments. Importantly, PhyloAln is designed to ideally remove common contaminations found in sequencing data. It is able to facilitate the streamlined preparation of sequence alignments for large-scale phylogenetic and evolutionary analyses in the current omic era.

## Materials and Methods

### Technological processes of PhyloAln

PhyloAln requires one or more reference alignments in FASTA format and one or more read/sequence files in FASTQ/FASTA format as input. The mapping of sequences into the alignments relies primarily on a search tool utilizing profile hidden Markov models, HMMER3 v3.1b2 (Mistry, et al. 2013), with preparation and downstream processes implemented using Python. To accommodate diverse data types, we have designed four search modes, namely ‘dna,’ ‘prot,’ ‘codon,’ and ‘dna_codon’. The ‘dna’ mode is tailored for direct HMMER3 search, suitable for nucleotide-to-nucleotide or protein-to-protein alignments, with an optional consideration for the reverse complement of sequences, particularly applicable in nucleotide-to-nucleotide alignments. The ‘prot’ mode and ‘codon’ mode necessitate protein alignments or codon alignments as references, respectively. In these modes, protein alignments directly provided or translated from codon alignments are employed to search six translation frames of the sequences using HMMER3. The ‘dna_codon’ mode shares similarities with the ‘dna’ mode in the HMMER3 search but additionally generates protein alignments by compulsively translating the output nucleotide alignments. This mode is suitable for codon-to-nucleotide alignment, for instance, when mapping long reads with a high ratio of insertions and deletions (Goodwin, et al. 2016) causing translation frame shifts into the alignments. All translations can be executed using standard or nonstandard genetic codes. Moreover, both the reference alignments and sequences can be optionally fragmented into smaller segments with a specified sliding length and step. This feature is designed for handling long alignments with time-consuming HMMER3 searches and genomic sequences with discontinuous coding regions, respectively. The HMMER3 search, combined with these preparations, facilitates the alignment of nuclear protein-coding genes, as well as protein-coding sequences with nonstandard genetic codes, non-protein-coding sequences, translated amino acid sequences, and various types of omic data.

Following the HMMER3 search, the matched sequences for each gene are subsequently extracted individually. Detailed information from the HMMER3 output files is then utilized to map the matched regions, including gaps, of the extracted sequences into corresponding coordinates within the reference alignments. In the ‘prot’ mode and ‘codon’ mode, the codons are associated with their respective amino acid sites. Optionally, these matched regions can be assembled into contigs based on overlapping coordinates, with specified overlapped length and percent identity thresholds.

To mitigate the impact of foreign contamination and poorly aligned regions, we have developed a conservative scoring method to assess the degree of matching for each target sequence in comparison to a specified outgroup in the reference alignments. Initially, the frequencies of each specific base at each site within the ingroup reference alignments (excluding the specified outgroup) are computed. Subsequently, for each target sequence, its conservative score is determined by the sum of the aforementioned frequencies of its bases at the respective sites. This score is then compared with the sum of frequencies of bases for the outgroup at the same coordinates. The conservative score of the outgroup can be adjusted by multiplying a weight coefficient, which is default as 0.9. If the conservative score of the target sequence is lower than the outgroup’s conservative score, the sequence is identified as foreign contamination and subsequently removed.

Multiple sources/species with their respective sequence files can be inputted through a configuration file. The optional detection of cross-contamination across the sources within the same configuration file is facilitated through sequence comparisons between any two sources, following processes similar to the methodology outlined in the CroCo software (Simion, et al. 2018). If the overlapped regions between two sequences surpass the specified length and percent identity, the two sequences are flagged as potential cross-contamination, and their abundance is scrutinized. The counts of reads constituting the two sequences are then normalized and compared. Given that reference alignments always do not encompass the entire gene sets, and standard transcript per kilobase per million mapped reads (TPM) and reads per kilobase per million mapped reads (RPKM) values cannot be directly calculated, we utilize an estimated value of RPKM to represent abundance. The estimated RPKM is computed by dividing the read counts by the sequence length in the reference coordinates and the total read count in the read files. If the fold of the estimated RPKM exceeds the specified threshold, the sequence with lower abundance is identified as cross-contamination from the source of the sequence with higher abundance and is subsequently removed.

Consensus of sites at the same reference coordinates and from the same sources can be optionally conducted before outputting the resultant alignments. Specifically, if the assembly step is disabled, cross-contamination will be detected and removed using the aforementioned method after the consensus of sequences. Ultimately, the aligned target sequences, with or without aligned reference sequences, are output to a new alignment FASTA file. In the ‘prot,’ ‘codon,’ and ‘dna_codon’ modes, translated protein alignments are also generated. All these steps are executed in parallel for computational efficiency.

### The simulated dataset

To assess PhyloAln’s capability in mapping sequences/reads into reference alignments across diverse species spanning the tree of life and its performance on contaminated data, we constructed a simulated dataset. This dataset comprises genomes from one bacterium (*Escherichia coli* (Hayashi, et al. 2001)), one plant (*Arabidopsis thaliana* (Cheng, et al. 2017)), one fungus (*Saccharomyces cerevisiae* (Engel, et al. 2022)), two vertebrates (*Danio rerio* (Howe, et al. 2013) and *Homo sapiens* (Collins, et al. 2004)), eight fruit flies (*Scaptodrosophila lebanonensis* (Flynn, et al. 2020), *Drosophila melanogaster* (Hoskins, et al. 2015), *D. simulans* (Wang, et al. 2023), *D. willistoni* (Ranz, et al. 2023), *D. mojavensis* (Kim, et al. 2021), *D. yakuba* (Huang, et al. 2022), *D. busckii* (Renschler, et al. 2019), *D. pseudoobscura* (Barata, et al. 2023)) and two other insects (*Tribolium castaneum* (Herndon, et al. 2020) and *Aedes aegypti* (Matthews, et al. 2018)), sourced from the NCBI RefSeq database (Haft, et al. 2023) (Table S1). The phylogeny of these species was obtained from the TimeTree database (Kumar, et al. 2022). Coding sequences, protein sequences, and transcripts were extracted based on General Feature Format (GFF) annotation files. For constructing reference alignments, *de novo* orthology assignment was performed using OrthoFinder v2.5.4 (Emms and Kelly 2019) on the longest transcript of the coding sequence for each gene in the aforementioned species. Subsequently, 46 single-copy genes among all the species were aligned using the L-INS-i mode of MAFFT v7.480 based on protein sequences (Katoh and Standley 2013). These alignments were then back-translated to codon alignments and trimmed using trimAl v1.4 (Capella-Gutierrez, et al. 2009) with the “automated1” option to eliminate poorly aligned regions. These trimmed alignments served as the reference alignments for evaluating the performance of PhyloAln.

The life-of-tree dataset comprises *E. coli*, *S. cerevisiae*, *D. rerio*, *T. castaneum* and *A. aegypti* as reference species, with *A. thaliana*, *H. sapiens* and *D. melanogaster* selected as target species. The five reference species were extracted from the 46 alignments, and sites with all gaps were removed to generate new reference alignment files for this dataset. The three target species were chosen to represent varying degrees of taxonomic relatedness to the closest reference species, spanning kingdoms (*A. thaliana*), classes (*H. sapiens*) and families (*D. melanogaster*). Simulated Illumina reads, with a length of 100 base pairs (bp), were generated using ART v2.5.8 (Huang, et al. 2012), while Nanopore and PacBio reads were simulated by ReadSim v1.6 (Lee, et al. 2014) with default average length. The coverage levels for genome and transcript sequences of the target species were set at 2X, 5X, 10X, 20X, and 30X, respectively.

For the contaminated dataset, transcripts from fruit fly species were mainly utilized, with *S. lebanonensis*, *D. mojavensis*, *D. yakuba*, *D. busckii*, *D. pseudoobscura* as reference species, and *D. melanogaster*, *D. simulans*, *D. willistoni* as target species. Additionally, *E. coli*, *A. thaliana*, *S. cerevisiae*, *D. rerio*, *H. sapiens*, *T. castaneum* and *A. aegypti* were included as foreign contamination sources. The new reference alignment files for the five reference species were prepared as described above. Illumina reads of the transcripts from target and foreign species were simulated by ART with a coverage of 10X, labeled with the source species for downstream testing. For each target species, reads were randomly selected 15,000,000-20,000,000 times. These reads were combined with 10,000-500,000 random reads from each foreign species and two other target species to introduce simulated foreign and cross-contamination, respectively. To evaluate the tools’ performance on sequences contaminated without being covered by large amounts of real target reads in the sequence consensus, five of the 46 single-copy genes in each target species were respectively selected to remove reads from the target regions while retaining the contamination reads.

### The real dataset of ladybird beetle transcriptomes

We selected genomes of 11 ladybird beetle (Coccinellidae) species (*Adalia bipunctata* (Wellcome Sanger Institute Tree of Life programme, et al. 2022), *Coccinella septempunctata* (Crowley, et al. 2021), *Cryptolaemus montrouzieri* (Li, Huang, et al. 2021), *Cynegetis impunctata* (our unpublished data, NCBI accession: GCA_030704885.1), *Halyzia sedecimguttata* (Crowley, et al. 2023), *Harmonia axyridis* (Boyes, et al. 2021), *Henosepilachna vigintioctomaculata* (Zhu, et al. 2023), *Henosepilachna vigintioctopunctata* (our unpublished data, NCBI accession: GCA_030704895.1), *Micraspis discolor* (our unpublished data, NCBI accession: GCA_030674115.1), *Novius pumilus* (Tang, et al. 2022) and *Propylea japonica* (Zhang, et al. 2020)), and one Coccinelloidea outgroup *Dastarcus helophoroides* (Zhang, et al. 2023), as reference species (Table S1). The genomes were downloaded from the NCBI Genome database. Gene predictions for *C. septempunctata* and *H. axyridis* were obtained from the NCBI RefSeq database, while those for the other ten genomes were generated using a modified FunAnnotate v1.8.1 pipeline (https://github.com/nextgenusfs/ funannotate), following the methodology outlined by Tang, et al. (2022).

The Illumina read files for 30 target ladybird beetle transcriptomes were obtained from the supplementary NCBI SRA accessions provided by Li, Tang, et al. (2021) (Table S1). The data were independently *de novo* assembled using Trinity v2.8.5 (Grabherr, et al. 2011). CroCo v1.1 (Simion, et al. 2018) was employed with default parameters to identify and eliminate cross-contamination sequences within the same batch. I Additionally, a customized nucleotide sequence database for Coccinelloidea was created, encompassing all ladybird genome sequences used in this study, alongside previously published versions (Ando, et al. 2018; Gautier, et al. 2018; Chen, et al. 2021) and the sequences of species under Coccinellidae, Endomychidae, Corylophidae, Latridiidae, Alexiidae, Cerylonidae, Discolomatidae, and Bothrideridae in the NCBI NT database. Subsequently, transcriptomic assemblies were queried against both this customized database and the NCBI NT database using BLAST v2.8.1+ (Camacho, et al. 2009). Transcripts with a percent identity < 90% compared to those within our customized database and a percent identity >= 98% compared to those in the NCBI NT database were deemed foreign and consequently removed from the assemblies. Non-redundant coding gene sets for each species were constructed using EvidentialGene v2018.06.18 (Gilbert 2013).

Orthology assignment, sequence alignment, and trimming were conducted across all 42 species as outlined for the simulated dataset. The 30 target species were then excluded from the alignments of a total of 1290 single-copy genes across 12 genomes. The resulting new alignments, devoid of all-gap sites, were utilized as reference alignments.

### The real dataset of pepper plastomes

For plant plastomes, we utilized the dataset from Simmonds, et al. (2021) to evaluate PhyloAln. The raw Illumina reads of 17 plastomes, along with the gene predictions and assemblies of all 36 plastomes (inclusive of the 17 target plastomes), were acquired from NCBI, based on the supplementary accessions or directly from the supplement materials (Table S1). Protein sequences of 68 single-copy protein-coding genes across all 36 plastomes were aligned using the L-INS-i mode of MAFFT and subsequently back-translated to codon alignments using the genetic code of 11 (the plant plastid code). The reference alignments, excluding the 17 target plastomes, were prepared as described earlier.

### The real dataset of turtle UCEs

The turtle UCE dataset from Crawford, et al. (2015) was employed to assess PhyloAln. The concatenated UCE matrix, encompassing a total of 38 species, was obtained from the supplementary materials. Raw Illumina reads for 27 target species were retrieved from the NCBI SRA database using the provided supplementary accessions (Table S1). Reference alignments, excluding the target sequences, were prepared as previously described.

### Run with PhyloAln, Read2Tree and Orthograph

In the simulated dataset spanning the tree of life, reads with varying coverages of genome and transcript sequences from *A. thaliana*, *H. sapiens* and *D. melanogaster* were aligned to the reference alignments using PhyloAln and Read2Tree v0.1.5 (Dylus, et al. 2024), respectively. *E. coli* was designated as the outgroup in PhyloAln. Due to the extensive memory requirements for the preparation of a large number of sequences in deep-depth read files on our computer server, especially for *H. sapiens*, a slower but less-memory-intensive preparation method, without storage of the sequences in the Python dictionary, was developed in PhyloAln and employed for these tests involving large-depth reads. For Illumina reads, the ‘codon’ mode was used in PhyloAln, while Read2Tree was run with default parameters. For PacBio and Nanopore long reads, the ‘dna_codon’ mode of PhyloAln was used to generate the alignments. When mapping the long reads of the genomes, the reads were split into fragments of 200 bp with a sliding step of 100 bp. Read2Tree for long reads also split the reads into fragments of 200 bp with ngmlr parameters for PacBio and Nanopore reads, respectively. Additionally, ‘codon’ mode without assembly of PhyloAln was used to align the complete sequences of the genome and transcript sequences to the reference alignments. Specifically, the genome sequences were initially split into fragments of 200 bp with a sliding step of 100 bp to mitigate the impact of intron regions.

Contaminated transcriptomic reads from *D. melanogaster*, *D. simulans*, *D. willistoni* were aligned to the reference alignments of fruit flies using both PhyloAln and Read2Tree. Specifically, for PhyloAln, ‘codon’ mode with and without assembly was employed, using *S. lebanonensis* as the outgroup.

In the real dataset of ladybird beetle transcriptomes, PhyloAln, Read2Tree, and Orthograph v0.7.2 (Petersen, et al. 2017) were employed. Due to the large number of reference alignments, PhyloAln was executed using ‘codon’ mode without assembly. Subsequently, reads and assemblies after decontamination were mapped into the reference alignments, respectively, with *D. helophoroides* set as the outgroup in PhyloAln. Read2Tree and Orthograph were run with default parameters, utilizing reads and decontaminated assemblies, respectively.

In addition, the performance of PhyloAln was evaluated using datasets of pepper plastomes and turtle UCEs. For pepper plastomes, assembly-free ‘codon’ mode with the genetic code of 11 was employed. Due to the presence of closely related and even the same species in the target list, making cross-contamination detection challenging (Simion, et al. 2018), the step of cross-contamination removal was disabled. *Magnolia grandiflora* was selected as the outgroup following the specifications of the pepper plastome dataset (Simmonds, et al. 2021). For the concatenated UCE matrix, the reference alignment was initially split into short alignments of 1,000 bp, and reads were then mapped into these short alignments using the ‘dna’ mode without assembly. Subsequently, all new short alignments were concatenated into the final matrix, with all processes executed within PhyloAln. *H. sapiens* was set as the outgroup based on the phylogenetic tree presented in the turtle UCE article (Crawford, et al. 2015).

PhyloAln utilized the prepared reference alignments directly in the dataset, while the unaligned sequences of each orthologous gene were provided to Read2Tree and Orthograph. All runs of PhyloAln, Read2Tree, and Orthograph used 20 threads, and their execution time was recorded using the ‘time’ command in the Linux system.

### Performance test of PhyloAln, Read2Tree and Orthograph

We calculated the completeness and percent identity of each sequence generated by PhyloAln, Read2Tree, and Orthograph in the alignments based on the reference coordinates. Only the sites with bases or intermediate gaps in the reference sequences and the generated sequences were considered valid regions, while the unknown or unmapped sites or those with start or end gaps were ignored. Completeness was defined as the percentage of overlapped valid regions in the valid regions of the reference sequences. Percent identity was defined as the percentage of the same site in both sequences among the overlapped valid regions. For PhyloAln, the generated sequences in the result alignments were compared with the corresponding sequences in the *de novo* assembled and aligned reference alignments before removal of the target sequences. Read2Tree generated the reference alignments itself in the pipeline, and we thus mapped the corresponding assembled sequences into these reference alignments using the ‘--add’ option and L-INS-i mode of MAFFT to obtain the reference templates. The sequences in the Read2Tree result alignments were then compared with the corresponding sequences in the reference templates. For Orthograph, since only the unaligned target sequences were obtained, we mapped the result sequences and the putative orthologous sequences into its intermediately generated reference alignments using the ‘--add’ option and L-INS-i mode of MAFFT, and compared them as described above. Specifically, in the dataset of ladybird transcriptomes, only 121 single-copy orthologous genes across all the 42 species, based on orthology assignments by OrthoFinder, were used for the test of completeness and percent identity. This was because these three tools only generated single-copy alignments in those 30 target species, making it challenging to compare with genes having multiple-copy sequences from the transcriptome assemblies in OrthoFinder results.

To assess the ability of PhyloAln to remove contamination, we utilized the species label of the reads and quantified the reads from each species composing the retained or removed sequences. We calculated the values of true positive (TP), true negative (TN), false positive (FP), false negative (FN), precision (= TP/(TP+FP)), and recall (= TP/(TP+FN)) for three aspects (clean, foreign contamination, cross contamination) in each target species.

To assess the ability of PhyloAln to remove contamination, we utilized the species label of the reads and quantified the reads from each species composing the retained or removed sequences. We calculated the values of true positive (TP), true negative (TN), false positive (FP), false negative (FN), precision (= TP/(TP+FP)), and recall (= TP/(TP+FN)) for three aspects (clean, foreign contamination, cross contamination) in each target species. The theoretical and observed positive/negative values were defined as follow:

#### Clean Aspect

The counts of reads from the target species were considered as theoretical positive values.

The counts of reads from other species were considered as theoretical negative values.

Reads retained in the final alignments were considered as observed positive values.

Reads removed were considered as observed negative values.

#### Foreign Contamination Aspect

The counts of reads from the foreign species were considered as theoretical positive values.

The counts of reads from the three target species were considered as theoretical negative values.

Reads removed in this step were considered as observed positive values.

Reads retained were considered as observed negative values.

#### Cross Contamination Aspect

The counts of reads from the other two target species were considered as theoretical positive values.

The counts of reads from the source species were considered as theoretical negative values.

Reads removed in this step were considered as observed positive values.

Reads retained were considered as observed negative values.

Due to relatively high false positive (FP) values in foreign contamination detection (putative foreign contamination from the target species), we additionally examined these putative contamination sequences. Alignments containing only these putative contamination sequences were output, and completeness and percent identity were calculated as described above. Protein and codon alignments of PhyloAln results and the putative contaminations were concatenated into a supermatrix and imported into the “MFP+MERGE” mode of IQ-TREE v2.1.4-beta (Minh, et al. 2020) with 1,000 ultrafast bootstrap (UFBoot) replicates, respectively.

In the three real datasets, final phylogenetic trees based on the results generated by PhyloAln, Read2Tree, and Orthograph were reconstructed using IQ-TREE with 1,000 ultrafast bootstrap (UFBoot) replicates. For Orthograph, the result sequences were initially aligned using the L-INS-i mode of MAFFT. The resulting alignments from the three tools were concatenated into a supermatrix and imported into the “MFP+MERGE” mode of IQ-TREE. Protein alignments were used in the ladybird transcriptome dataset, while codon alignments were employed in the pepper plastome dataset. In the UCE matrix dataset without partition information, the concatenated alignment output by PhyloAln was utilized for phylogenetic analysis by IQ-TREE with 1,000 UFBoot replicates and default parameters. Reference trees were constructed based on concatenated protein supermatrices comprising 1290 single-copy genes from OrthoFinder results in the ladybird transcriptome dataset, concatenated codon supermatrices consisting of 68 plastid protein-coding genes from pepper plastome assemblies, and the downloaded UCE matrix. All the trees were rooted with the corresponding outgroup mentioned in PhyloAln processes using the Python ETE 3 Toolkit (Huerta-Cepas, et al. 2016) and visualized with the R package ggtree (Yu, et al. 2017).

### Check of the source of *Graptemys pseudogeographica* reads in the turtle UCE dataset

Due to the different positions of *Graptemys pseudogeographica* between the phylogenetic trees obtained from PhyloAln results using read files and the downloaded sequence matrix, along with the relatively low completeness and percent identity of the *G. pseudogeographica* sequence in the test, we investigated the source of its reads. We employed IDBA-UD v1.1.3 (Peng, et al. 2012) to assemble the reads, and the resulting assembly was used to search for sequences of all the species in the downloaded sequence matrix using BLAST v2.8.1+ (Camacho, et al. 2009) with an E-value threshold of 1e-5. The species with the best hits for each sequence in the assembly were then counted.

## Data and Resource Availability

References and all accessions of the genomes and reads used are listed in Table S1 of Supplementary File 2. The source code for PhyloAln is available under an MIT open-source license at https://github.com/huangyh45/PhyloAln.

## Authors’ contributions

YHH and HP designed the study. YHH implemented the software. YHH, HL and YFS performed data analysis, code review and software test. YHH, HSL and HP drafted the manuscript. All authors read and approved the final version of the manuscript.

## Acknowledgements

This work was supported by the National Natural Science Foundation of China (Grant No. 32172472, 31970439), Open Fund of Guangdong Key Laboratory of Animal Protection and Resource Utilization (Grant No. GIZ-KE202304) and National Key R&D Program of China (Grant No. 2023YFD1400600).

