## Supplementary Figures S1-S11. for "PhyloAln: a convenient reference-based tool to align sequences and high-throughput reads for phylogeny and evolution in the omic era"

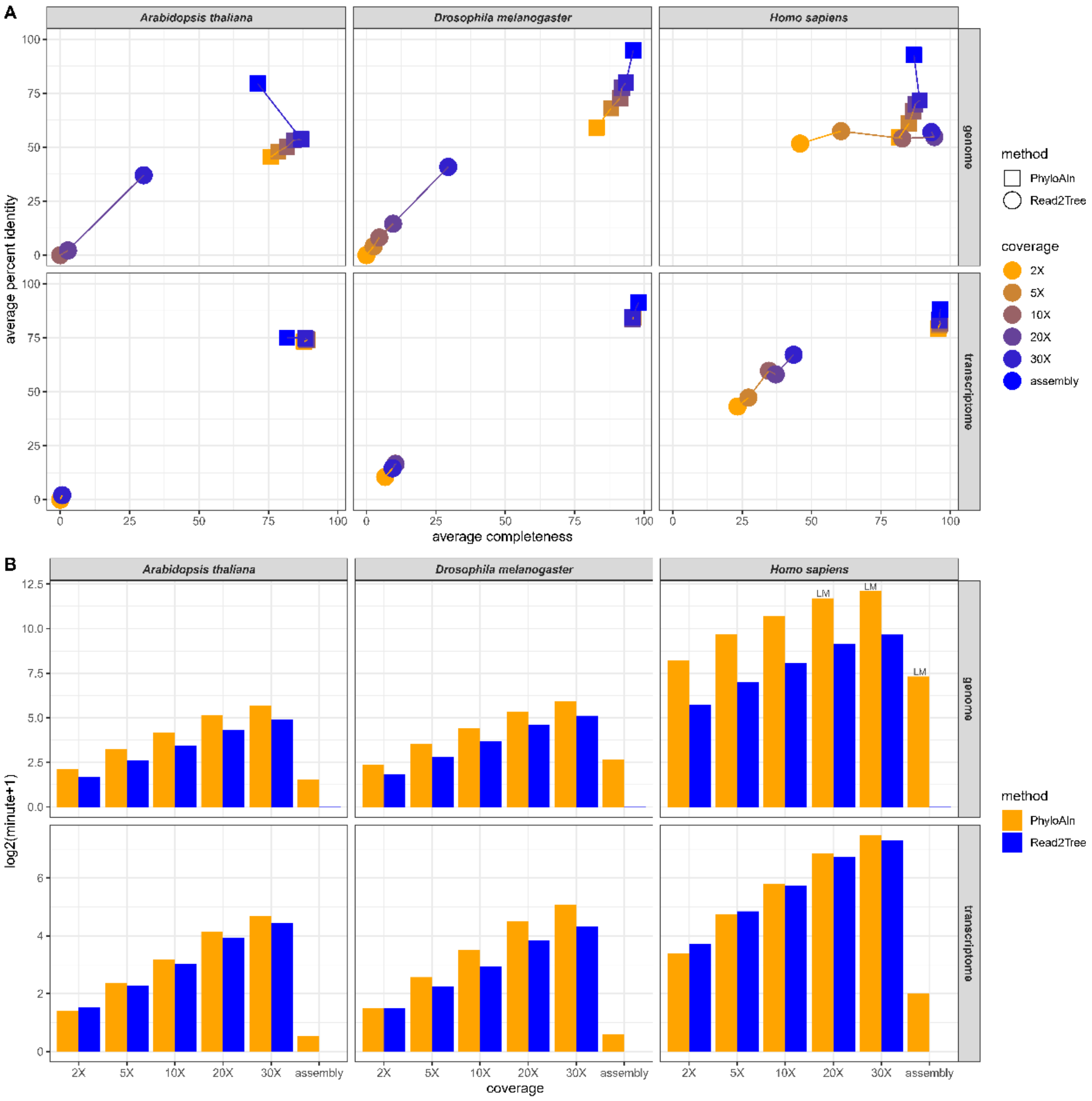


Figure S1. Performance test of PhyloAln and Read2Tree using PacBio reads on the simulated dataset across the tree of life. A) Average completeness and percent identity of the alignments generated by PhyloAln and Read2Tree using the simulated PacBio reads or original assemblies of genomes and transcriptomes of three target species with different coverages. C) Running time of PhyloAln and Read2Tree using the simulated PacBio reads or original assemblies of genomes and transcriptomes of three target species with different coverages. LM: using a slower low-memory strategy to prepare the sequences and reads because of too large size of data.

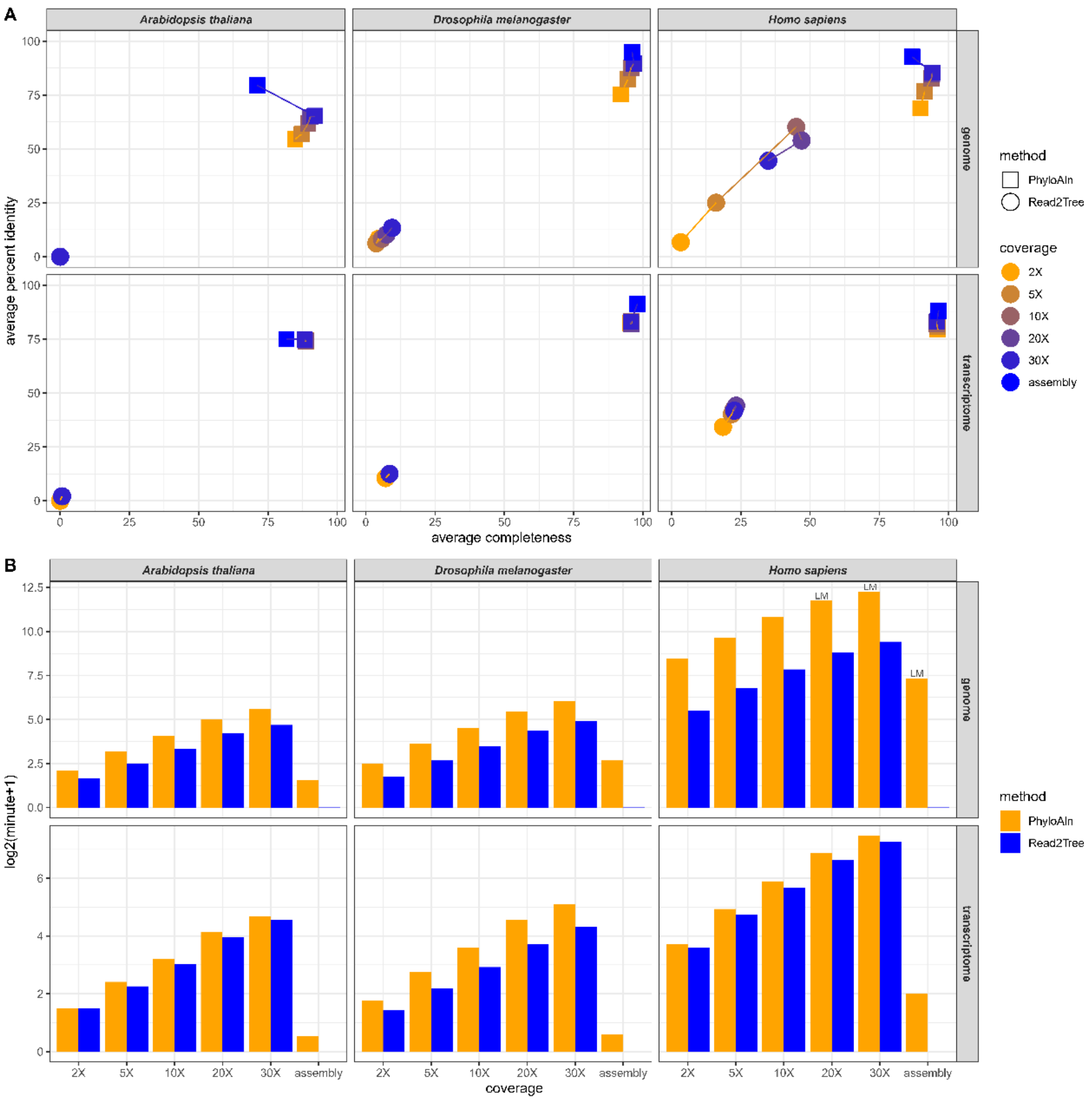


Figure S2. Performance test of PhyloAln and Read2Tree using Nanopore reads on the simulated dataset across the tree of life. A) Average completeness and percent identity of the alignments generated by PhyloAln and Read2Tree using the simulated Nanopore reads or original assemblies of genomes and transcriptomes of three target species with different coverages. C) Running time of PhyloAln and Read2Tree using the simulated Nanopore reads or original assemblies of genomes and transcriptomes of three target species with different coverages. LM: using a slower low-memory strategy to prepare the sequences and reads because of too large size of data.

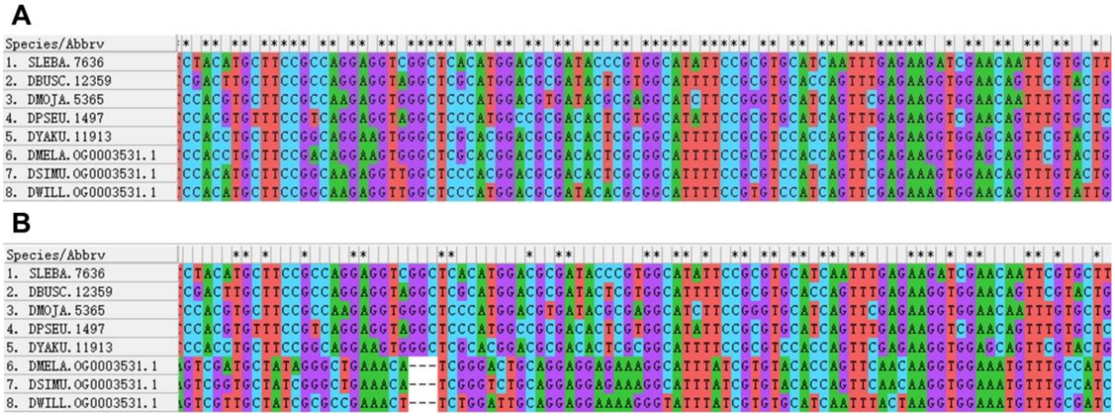

Figure S3. Poor alignments of the consensus sequences of the reads from the target species detected as foreign contamination. A) The PhyloAln result alignments. B) The alignments of the consensus sequences of the reads from the target species detected as foreign contamination.

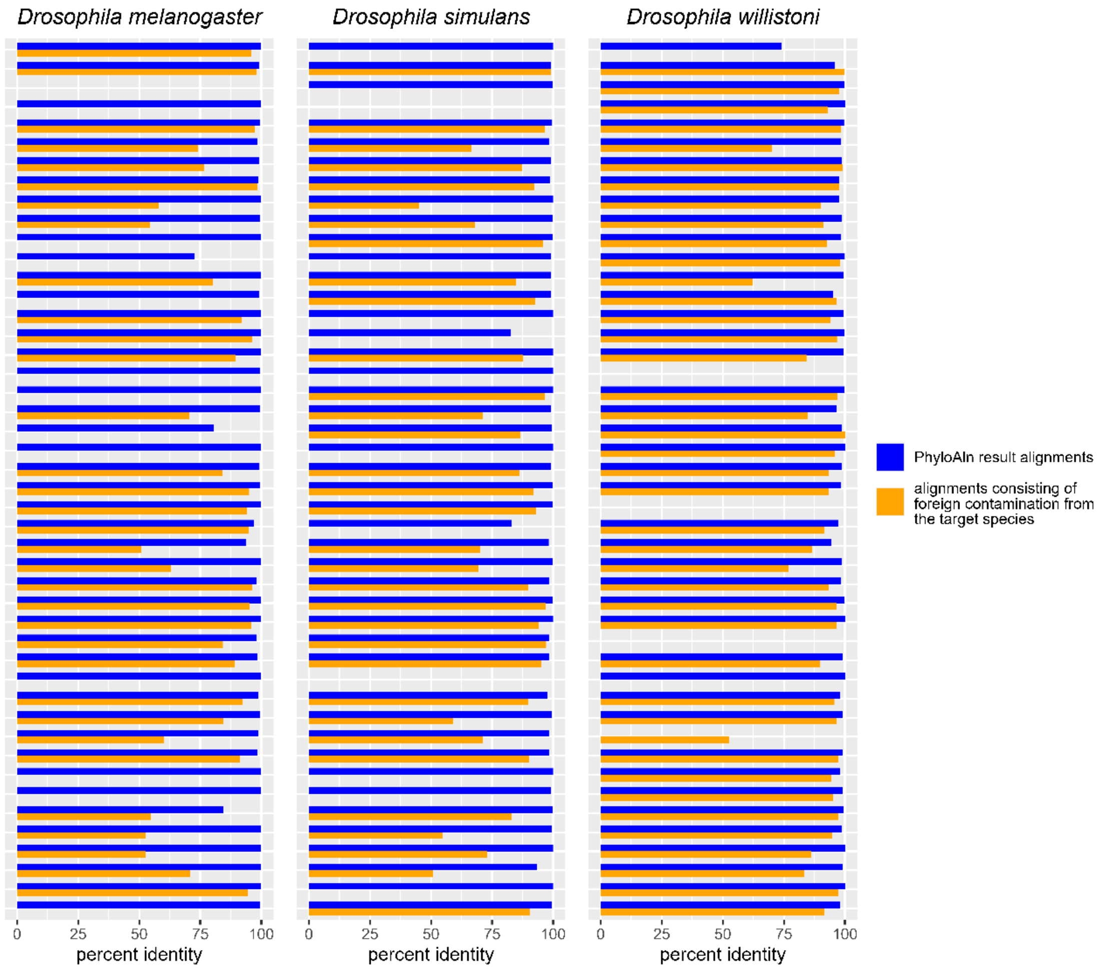


Figure S4. Percent identity of each orthologous gene of the PhyloAln result alignments and the alignments of the consensus sequences of the reads from the target species detected as foreign contamination.

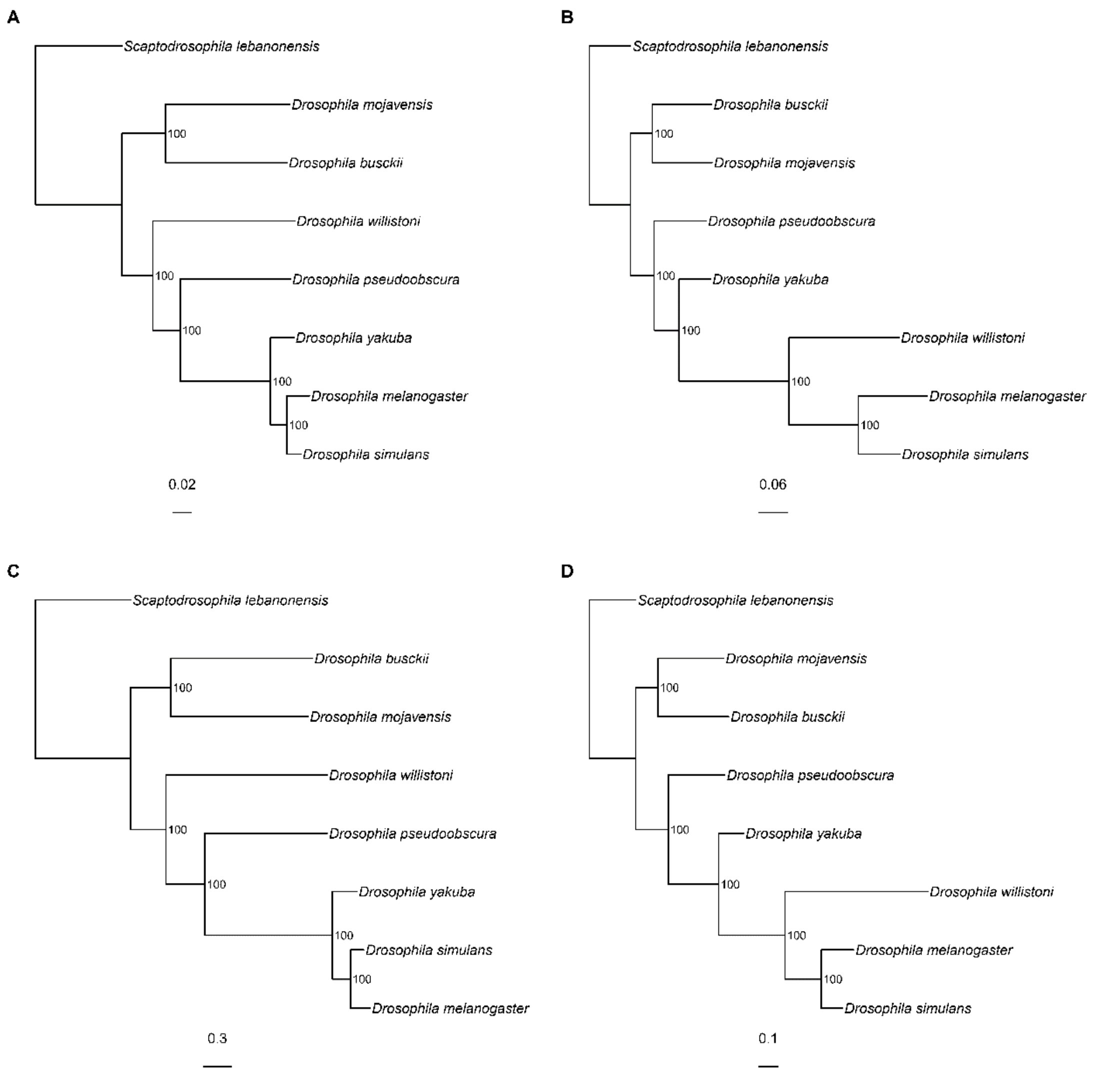


Figure S5. Incorrect phylogeny based on the alignments of the consensus sequences of the reads from the target species detected as foreign contamination. A) Phylogenetic tree reconstructed by IQ-TREE based on the PhyloAln result protein alignments. B) Phylogenetic tree reconstructed by IQ-TREE based on the protein alignments translated from the consensus sequences of the reads from the target species detected as foreign contamination. C) Phylogenetic tree reconstructed by IQ-TREE based on the PhyloAln result codon alignments. D) Phylogenetic tree reconstructed by IQ-TREE based on the codon alignments of the consensus sequences of the reads from the target species detected as foreign contamination. The numbers beside the nodes are the ultrafast bootstrap (UFBoot) support values obtained by IQ-TREE.

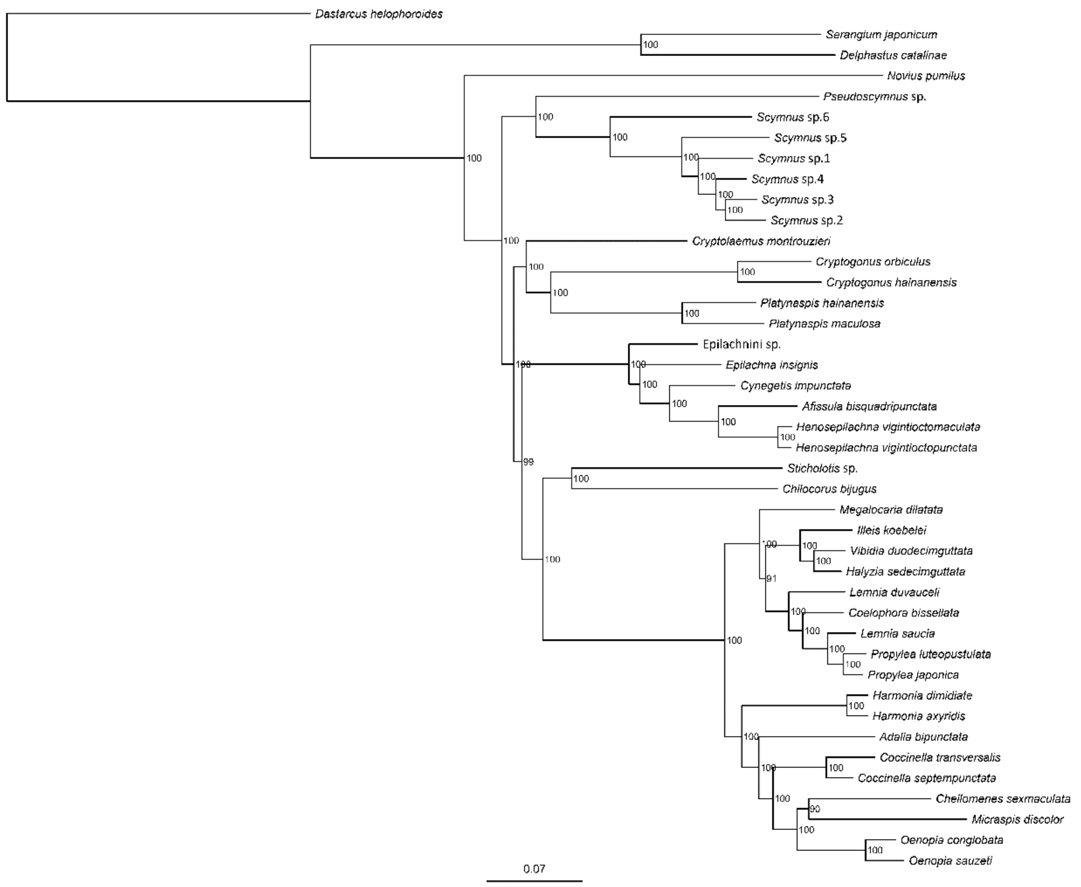


Figure S6. Phylogenetic tree reconstructed by IQ-TREE based on the protein alignments generated by OrthoFinder and MAFFT using 12 reference Coccinelloidea genomes and 30 target ladybird beetle (Coccinellidae) transcriptomes. The numbers beside the nodes are the ultrafast bootstrap (UFBoot) support values obtained by IQ-TREE.

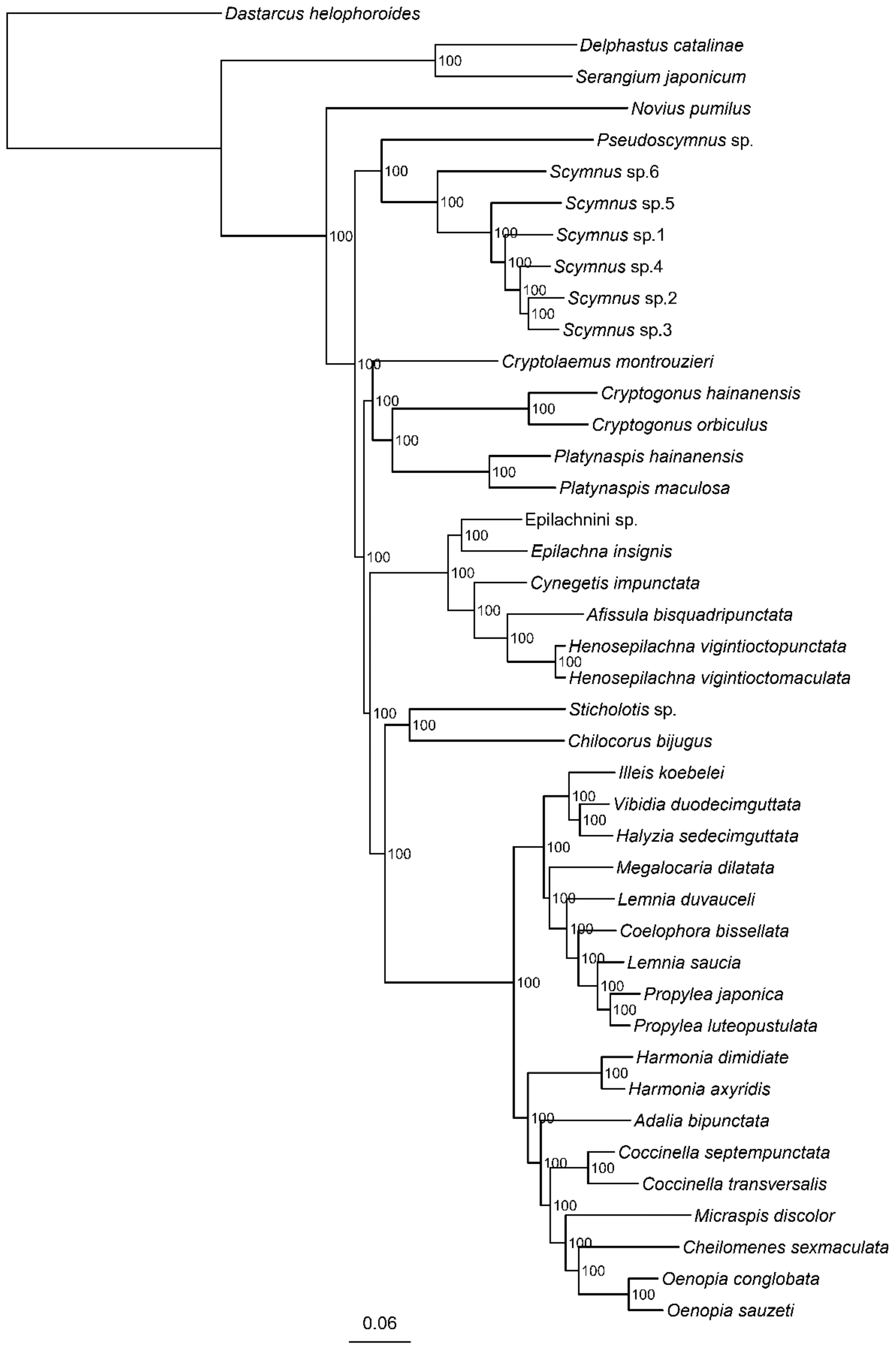


Figure S7. Phylogenetic tree reconstructed by IQ-TREE based on the protein alignments generated by PhyloAln using 12 reference Coccinelloidea genomes and the reads of 30 target ladybird beetle (Coccinellidae) transcriptomes. The numbers beside the nodes are the ultrafast bootstrap (UFBoot) support values obtained by IQ-TREE.

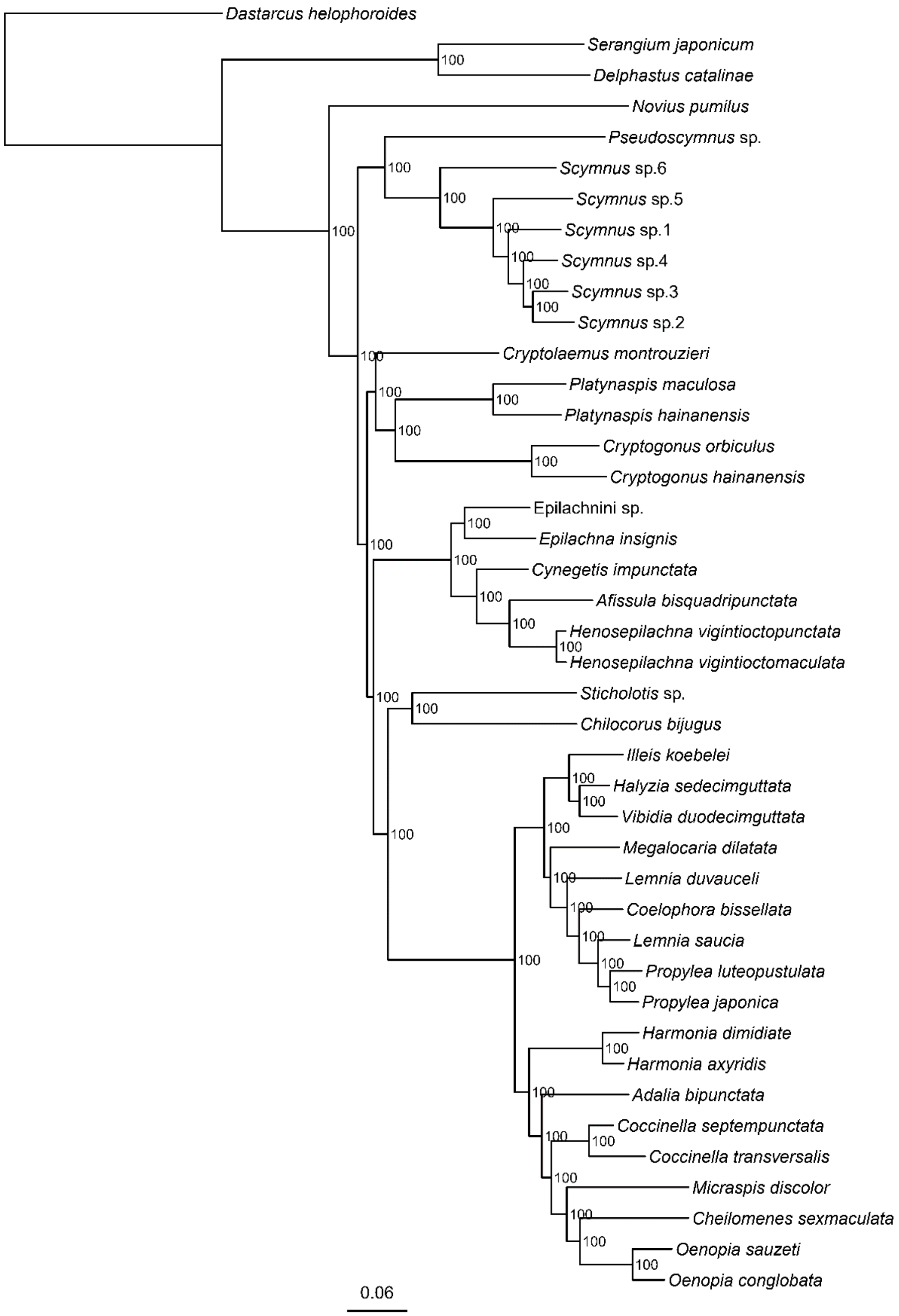


Figure S8. Phylogenetic tree reconstructed by IQ-TREE based on the protein alignments generated by PhyloAln using 12 reference Coccinelloidea genomes and the decontaminated assemblies of 30 target ladybird beetle (Coccinellidae) transcriptomes. The numbers beside the nodes are the ultrafast bootstrap (UFBoot) support values obtained by IQ-TREE.

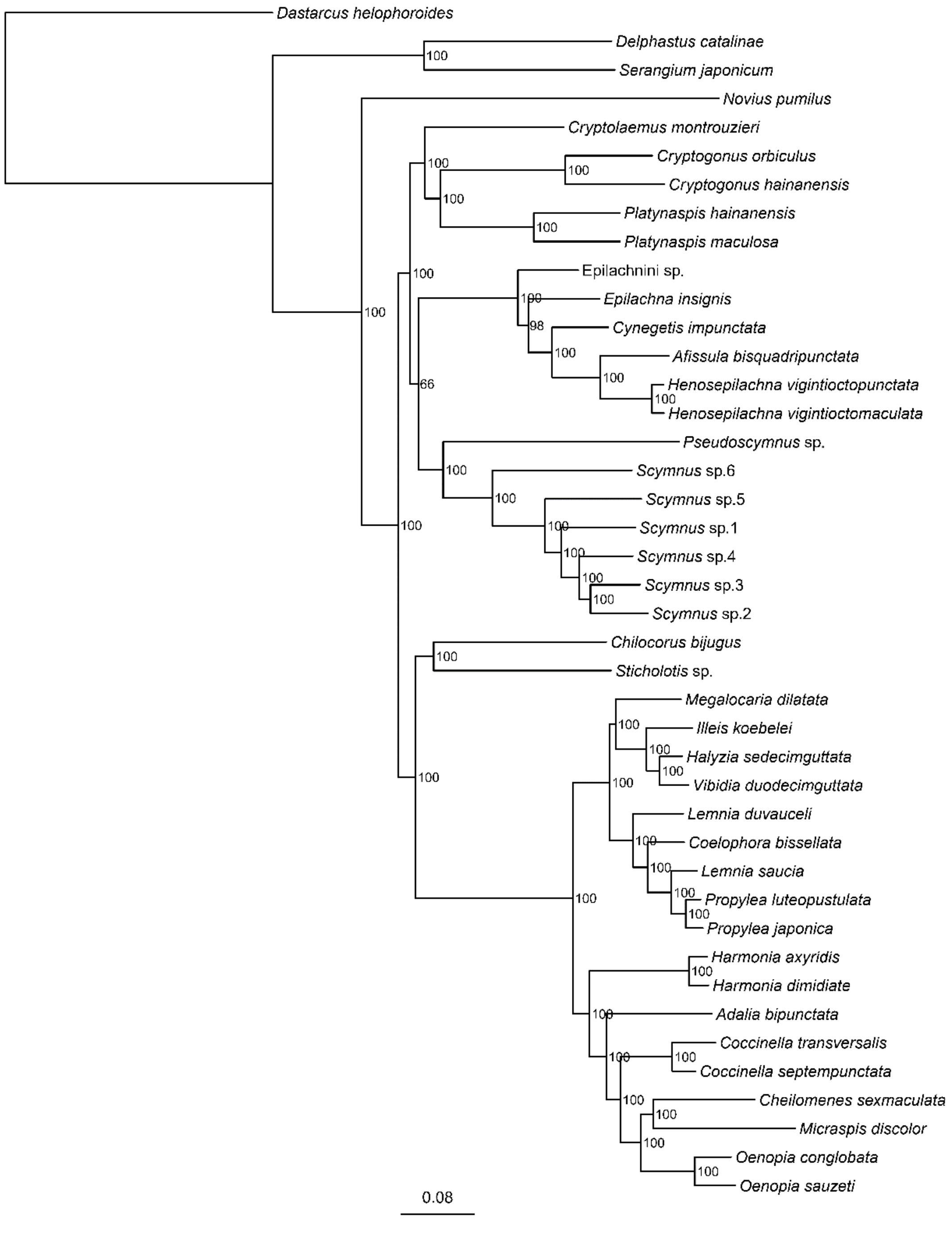


Figure S9. Phylogenetic tree reconstructed by IQ-TREE based on the protein alignments generated by Read2Tree using 12 reference Coccinelloidea genomes and the reads of 30 target ladybird beetle (Coccinellidae) transcriptomes. The numbers beside the nodes are the ultrafast bootstrap (UFBoot) support values obtained by IQ-TREE.

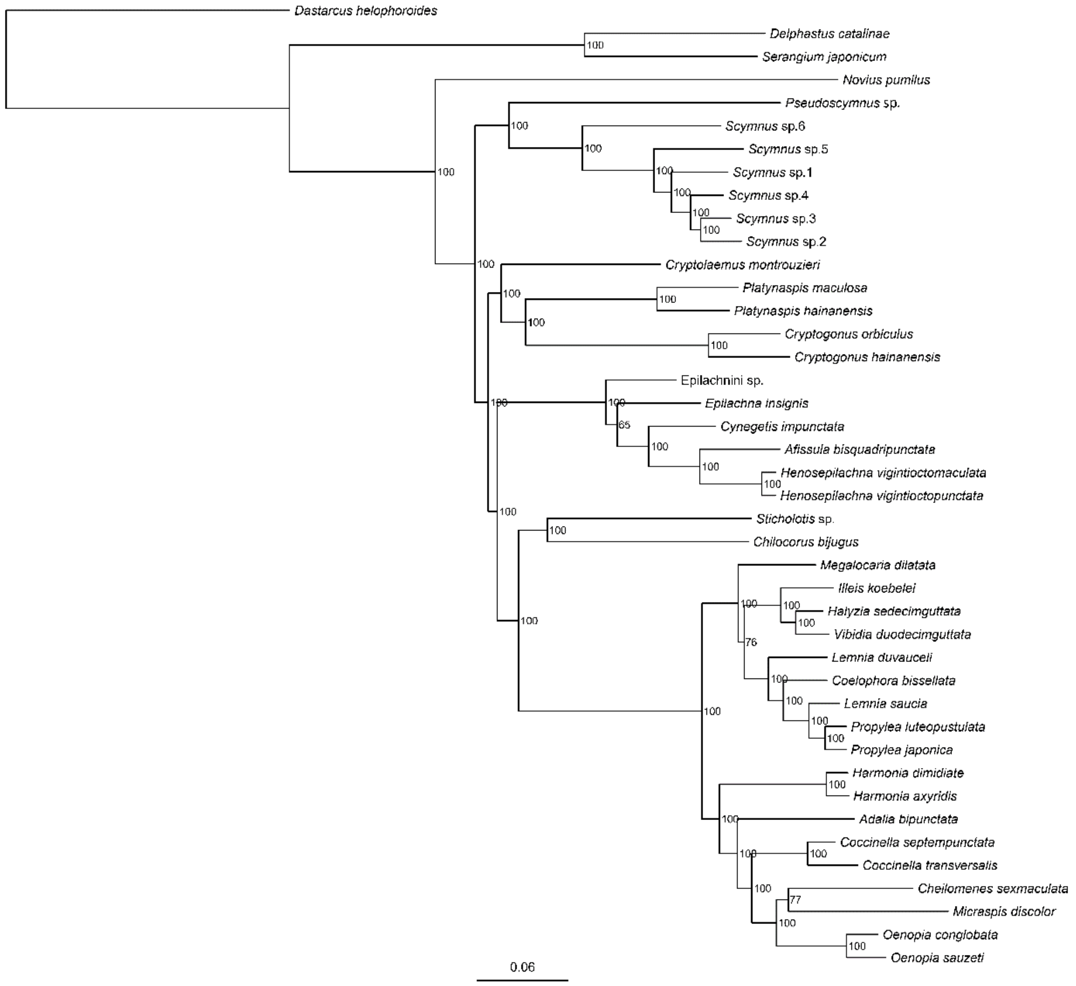


Figure S10. Phylogenetic tree reconstructed by IQ-TREE based on the protein alignments generated by Orthograph using 12 reference Coccinelloidea genomes and the decontaminated assemblies of 30 target ladybird beetle (Coccinellidae) transcriptomes. The numbers beside the nodes are the ultrafast bootstrap (UFBoot) support values obtained by IQ-TREE.

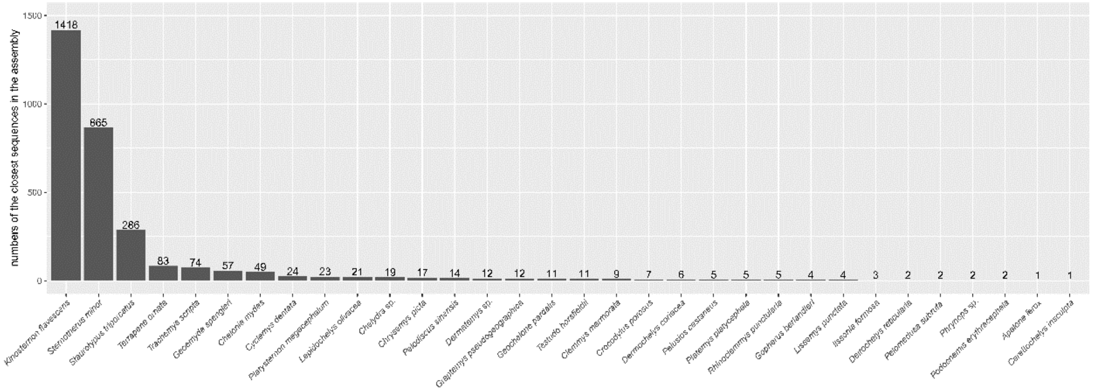


Figure S11. Numbers of the assembled sequences from the reads of *Graptemys pseudogeographica* closest to each sequence in the supplementary concatenated turtle ultraconserved element (UCE) matrix of Crawford, et al. (2015).
